# Can evolutionary potential help plants survive climate change? High variability in evolvability and G-matrix structure of *Hypericum* populations

**DOI:** 10.1101/2025.11.10.687560

**Authors:** Anniina L. K. Mattila, Marko-Tapio Hyvärinen, Maria H. Hällfors, Susanna H. M. Koivusaari, Charlotte Møller, Laura Pietikäinen, Øystein H. Opedal

## Abstract

To predict the consequences of global change, we need knowledge of the ability of populations to evolve. On the short term, this is driven by the amount of standing genetic variation in the population. There is a pressing need for empirical data on evolutionary potential of natural populations, and on how evolutionary potential varies among species and across species distributions. Here, we evaluate and compare additive genetic variance-covariance matrices (G-matrices) and evolutionary potential in multiple populations of two *Hypericum* species, one of them a generalist perennial plant with a wide geographic distribution, and the other a rare habitat specialist. With these comparisons, we aim to evaluate the stability of the G-matrix within and among plant species, and whether the genetic architecture of some plant trait modules remains more stable than of others. We utilize manual crossing experiments in a nested half-sibling breeding design and quantitative genetic analyses to estimate evolvabilities (mean-scaled genetic variances) as a proportional measure of evolutionary potential. With such comparisons of evolvability in different types of species and populations, we can begin to understand general patterns in the variation of evolutionary potential in nature.

## 3. INTRODUCTION

To accurately predict the capacity of natural populations to adapt to ongoing and future global changes, precise estimates of evolutionary potential are urgently needed. Environmental change often leads to changes in the adaptive optimum phenotype, and hence in the strength and mode of natural selection. The ability of populations to adapt through response to this selection depends on the amount of standing genetic variation in the traits under selection. Furthermore, the inheritance of a trait is seldom independent of other traits of the organism. In fact, selection and adaptation are multivariate processes, and the response to selection also depends on the standing genetic covariances among the evolving traits. In evolutionary quantitative genetics, phenotypic variances and covariances are summarized in compact form in variance matrices comprising variances on the diagonal and covariances elsewhere (Lande, 1979; Hansen and Houle, 2008). To forecast response to multivariate selection, the variance matrix for additive-genetic (breeding) values (G) is of special interest, because it combines with a multivariate selection gradient to yield the expected magnitude of the selection response (Lande and Arnold, 1983).

Recent work has demonstrated the evolutionary relevance of the G-matrix by showing that G estimated for a single population often aligns surprisingly well with patterns of population divergence (Schluter, 1996; Houle *et al*., 2017; McGlothlin *et al*., 2018; Opedal *et al*., 2023). This indirectly suggests that the structure of G is rather stable across conspecific populations, because substantial variation in G would make it highly unlikely that an estimate for a randomly chosen population would predict patterns of divergence. However, there are also reasons to believe that G can evolve during the diversification process, potentially so that it comes into better alignment with patterns of divergence (Arnold *et al*., 2008; Arnold, 2023).

Different kinds of traits are known to differ both in the amount of additive genetic variance and in patterns of genetic covariances. In plants, flowers are known to be integrated structures (i.e. the phenotypes of different structures within the flower are co-dependent) and to exhibit less additive genetic variance than do vegetative traits, presumably as a result of pollinator-mediated stabilizing selection (Opedal, 2019). Vegetative traits are also known to be integrated, and to vary considerably among environments. In contrast, little is known about the stability of floral vs. vegetative G-matrices across populations and along environmental gradients, partly because floral and vegetative traits tend to be studied separately. Knowledge about the potentially differing evolutionary patterns and mechanisms of floral vs. vegetative traits has relevance for predicting evolutionary responses of plants in a changing world. For instance, the potential to evolve in response to changing water availability through a change in the vegetative phenotype may differ substantially from the potential for the floral phenotype to evolve to a changing pollinator environment.

Previous studies assessing variation in G-matrices among conspecific plant populations have yielded somewhat conflicting results, with some authors reporting apparent stability (Puentes, Granath and Ågren, 2016; Henry and Stinchcombe, 2023), while others have detected substantial variation (e.g.(Walter *et al*., 2018)). There are too few studies for general patterns to emerge. Patterns of variation in G-matrices thus remains unresolved in general, and for plants in particular. Specifically, whether floral vs. vegetative traits differ in patterns of variation in multivariate genetic architecture has not been explored.

Quantifying geographic variation in multivariate genetic architecture is also highly relevant for efforts towards predicting population responses to environmental change. To forecast future evolution, we need to know whether G-matrices estimated in single populations can be extrapolated to species level, and whether we can rely on G-matrices estimated in contemporary populations, or whether we also need to forecast future changes in G.

Here we explore the multivariate genetic architecture of floral, vegetative and life-history traits in the widespread Eurasian herb *Hypericum maculatum* and its less widespread congener *H. montanum*. We estimated G-matrices for three populations of *H. maculatum* and two populations of *H. montanum* and analyze these to explore the following questions. **1.** How variable is G-matrix size (i.e. evolvability) and structure across populations and species. **2.** Do these patterns of variation differ among G-matrices of different trait classes, e.g. floral and vegetative traits? **3.** How can the data and potential emerging patterns be applied for evaluating plant adaptive responses to climate change?

## 4. MATERIAL AND METHODS

### Study species and populations

The two studied *Hypericum* (*Hypericaceae*) species are perennial herbs native to Europe. *H. maculatum Crantz* is a habitat generalist species found on young meadows, pastures, forest margins and road-sides, and is prevalent within its European distributional range, while *H. montanum L.* is a sparsely-distributed calciphile habitat specialist of light forest margins and precipices. The northern range edge of both species is located in south-central Fennoscandia. Both species have unassisted dispersal with a maximum seed dispersal distance of 5 meters for 99 % of the seeds (class 2; (Lososová *et al*., 2023)). We acquired seeds from three *H. maculatum* and two *H. montanum* populations by collecting fresh seeds from the field in 2020-2021. The seeds were collected in paper bags separately per maternal individual, cleaned and stored in dry conditions in room temperature. The collection of fresh seeds of *H. montanum L.* in Finland, where the species is protected, was approved by the Centre for Economic Development, Transport and the Environment (UUDELY/8787/2021).

### Plant cultivation and crossing design

#### Parental generation

The seeds were cold stratified at 4°C for eight weeks before sowing. The cultivation and crossings of the parental generation individuals was undertaken in two batches: batch 1) seeds of populations Hmac_MHH5, Hmac_MH1 and Hmon_G were sown on Feb 25-26 2021, and batch 2) seeds of populations Hmac_Mei and Hmon_Loh on Jul 5-6 2022. From each population, we sowed 10 seeds per mother plant, either individually on plug trays in a 5:2:1.5 mixture of seedling peat (Kekkilä Professional W HS R8017), vermiculite and sand in greenhouse conditions set at 20/10°C 16h light / 8h dark at the Viikki Plant Growth Facilities (University of Helsinki) (batch 1), or five seeds per pot in two 10 x 10cm pots on seedling peat in semi-open greenhouse conditions at the Kumpula botanic garden (University of Helsinki) (batch 2). After sowing, the soil was covered with 0-2 mm sand and kept moist with automated or manual watering every 24-48 hours and water-retaining rugs. Two months after sowing, five randomly chosen seedlings per maternal family were transplanted into 1 L pots in 5:2:1 mixture of peat soil (Kekkilä Professional C1 R8089), vermiculate and sand (batch 1) or in 10:1 mixture of sandy peat soil (Kekkilä Professional W R8014) and vermiculite (batch 2), and kept them in a greenhouse (Viikki Plant Growth Facilities, University of Helsinki) at weekly rotated locations at 20/10°C 18/6h light/dark (batch 1) or 24/16°C 18/6h light/dark (batch 2). Because of soil compaction, batch 1 plants were further repotted five months after sowing into 1 L pots in 10:1 mixture of sandy peat soil (Kekkilä Professional W R8014) and vermiculite. The plants were automatically watered every 48h, and fertilized bimonthly with a 0.075-0.2% fertilizer solution (Kekkilä Professional Superex NPK 12-5-27). The plants were supported with bamboo sticks. Pest control and fungal infection treatment was administered as needed. Upon the development of the first buds, the plants from batch 1 were covered with plastic pollination bags, and the flowers from batch 2 were covered with nylon tulle bags to prevent uncontrolled pollination.

#### Crossing design

We performed crossings within each population, obtaining in total 314 full-sibling families (167 and 147 *H. maculatum* and *H. montanum* families, respectively; Table 1). The crosses were done manually, such that one designated sire from each maternal family (61 *H. maculatum* and 50 *H. montanum* sires) was crossed with randomly chosen dams (mean=2.5 and 3.0 dams per *H. maculatum* and *H. montanum* sire, respectively) from different maternal families within the same population. These crosses resulted in a nested full-sibling (offspring of each dam)/half-sibling (offspring of each sire) pedigree structure in the F1 generation. Several flowers of each dam were manually pollinated with pollen from the chosen sire. Before pollination, the flowers were emasculated (as late buds or freshly-opened flowers) to prevent self-pollination, by using dissection scissors to cut off all anthers while leaving stigmas intact. The emasculated and manually pollinated flowers were marked with a tied string. After all crosses were done, batch 1 plants were moved to cooler greenhouse conditions (12/8°C, natural light) to await fruit ripening. Batch 2 plants remained in the same greenhouse conditions until fruits ripened. Ripe seeds from the marked pollinated flowers (F1 generation seeds) were collected by dam in paper bags, cleaned and stored in dry room temperature.

**Table 1.** The study populations and their localities. (*exact location cannot be disclosed due to protection status), as well as species- and population-specific sample sizes of dataset 1 (includes data of traits: flower corolla diameter, stigma length, anther length, plant height, leaf length, number of leaf pairs, flower count and flower phenology) and dataset 2 (includes data of traits: plant height, leaf length, number of leaf pairs).

| Species | Population | Country | Lat | Lon | Dataset 1 |  | Dataset 2 |  |
| --- | --- | --- | --- | --- | --- | --- | --- | --- |
|  |  |  |  |  | n(F1) | n(sires) | n(F1) | n(sires) |
| *H. maculatum* | Hmac_pooled | Multiple | Multiple | Multiple | 256 | 58 | 508 | 61 |
| *H. maculatum* | Hmac_MHH5 | Finland | 60.18 | 24.70 | 87 | 23 | 220 | 25 |
| *H. maculatum* | Hmac_MH1 | Finland | 60.26 | 23.60 | 98 | 25 | 176 | 26 |
| *H. maculatum* | Hmac_Mei | Belgium | 50.50 | 6.25 | 71 | 10 | 112 | 10 |
| *H. montanum* | Hmon_pooled | Multiple | Multiple | Multiple | 68 | 20 | 337 | 50 |
| *H. montanum* | Hmon_G | Sweden | 57.79 | 18.89 | 68 | 20 | 181 | 22 |
| *H. montanum* | Hmon_Loh | Finland | 60* | 24* | NA | NA | 156 | 28 |

#### F1 generation

The F1 seeds were cold stratified at 4°C for eight weeks. We then sowed 10 seeds per maternal family, with five seeds per pot in two 10 x 10cm pots on seedling peat (Kekkilä Professional W HS R8017) in semi-open greenhouse conditions at the Kumpula botanic garden (University of Helsinki) on July 5-6 2022 (batch 1) or Aug 16 2023 (batch 2). After sowing, the soil was covered with 0-2 mm sand and kept moist with watering every 24-48 hours and water-retaining rugs. The position of pots was rotated bimonthly. Two months after sowing, 3 randomly chosen seedlings per maternal family were repotted into 1 L pots in 10:1 mixture of sandy peat soil (Kekkilä Professional W R8014) and vermiculite, and moved 2-3 months after sowing to a greenhouse (Viikki Plant Growth Facilities, University of Helsinki), where they were kept on water retaining rugs in biweekly rotated locations at 24/16°C 16/8h light/dark (batch 1 populations Hmac_MHH5 & Hmac_MH1), or 20/12°C 16/8h light/dark (batch 1 Hmon_G & batch 2 Hmac_Meise & Hmon_Loh). The plants were supported with bamboo sticks, watered 2-4 times weekly and fertilized bimonthly with a 0.2% fertilizer solution (Kekkilä Professional Superex NPK 12-5-27). Pest control and fungal infection treatment was administered as needed.

### Trait measurements

The F1 individuals were monitored 1-3 times weekly for the onset of flowering, and the date of the first open flower was recorded. At first flowering, we also recorded floral trait measurements from the topmost open flower, including 1) flower corolla diameter, 2) stigma length, and 3) anther length. Vegetative traits and flower count were recorded around the time of population peak flowering, simultaneously for all individuals in the population. Vegetative trait measurements included 1) plant height from soil level to the tip of tallest branch, 2) leaf length of the largest leaf, and 3) the number of leaf pairs (number of leaf nodes on the tallest branch). The total count of flowers included buds, open and wilted flowers and fruit. A digital caliper (Mitutoyo Absolute AOS DIGIMATIC Model CD-15APX, 0.01mm precision) was used for leaf and floral measurements. Plant height was measured with the help of a cotton string to track the length of the tallest branch, thereafter measuring the length of the string with a tape measure. *H. montanum* population Hmon_Loh did not proceed to flowering stage during the experiment, and we thus recorded only vegetative trait measurements for this population.

We divided the resulting trait data into two datasets (see Table 1). Dataset 1 consists of offspring individuals of 58 *H. maculatum* and 20 *H. montanum* (only population Hmon_G) sires for which we obtained measurements of both vegetative and floral traits: on average 4.9 and 3.4 offspring individuals of each *H. maculatum* and *H. montanum* sire, respectively, which resulted in data from a total of 256 *H. maculatum* and 68 *H. montanum* F1-generation individuals. Dataset 2 includes all individuals with vegetative trait measurements, obtained from on average 8.9 offspring individuals of each *H. maculatum* sire (61 sires in total) and 6.9 offspring individuals of each *H. montanum* sire (50 sires in total), resulting in data from a total of 508 *H. maculatum* and 337 *H. montanum* F1-generation individuals.

### Data analysis

#### Covariance of measured traits

To study covariance of traits in the raw F1 data, i.e. ignoring family relationships, we estimated a mean-scaled phenotypic variance matrix (*P*) by computing a covariance matrix of the studied traits, and calculating both population-specific *P*, as well as mean *P* over populations (separately for species). To estimate the relationship of the recorded traits with reproductive fitness, we additionally calculated the correlation of traits (Pearson correlation) with the expected closest proxies of reproductive output available: the total number of flowers (expected to correlate with pollen yield and fruit set; see e.g. Devlin, Clegg and Ellstrand, 1992; Opedal, 2021) and the average number of seeds per fruit, separately for each species.

#### Quantitative-genetic analysis

##### Estimating G-matrices

Because only a subset of the F1 individuals flowered during the data-collection periods, we analyzed two different subsets of the data. The first subset (Dataset 1) included individuals for which all traits (floral + vegetative) were measured, and the second and larger subset (Dataset 2) included all individuals for which the vegetative traits were measured. Using these datasets, we estimated three different genetic variance-covariance matrices (*G*). ***G_F+V_*** included floral and vegetative traits (flower corolla diameter, anther length, stigma length, plant height, leaf length and number of leaf pairs) from Dataset 1. ***G_F+V_*** was also further analyzed as sub-matrices consisting of only the floral or the vegetative traits. ***G_V_*** used the larger Dataset 2 and included the vegetative traits plant height, leaf length and number of leaf pairs. Flowering (yes/no) was included in the model for ***G_V_*** as a fixed factor. ***G_LH_*** consisted of a set of life-history and fitness traits (flower corolla diameter, plant height, number of leaf pairs, total flower count (log-transformed) and flowering phenology (days from sowing to opening of first flower) using Dataset 1.

To estimate the G-matrix (*G*), we fit a multivariate animal model with the MCMCglmm R package (Hadfield, 2010) of the form

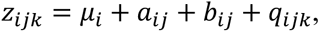

where *z* is the phenotypic trait value, *μ* is the trait mean, *a* is the breeding value, *b* is the non-genetic plant-level effect and *q* is the residuals. The indices run over traits (*i*), plants (*j*), and unique data points (*k*). We estimated the G-matrices separately for each population. We also estimated species-specific mean G-matrices by running the animal model with pooled data for each species, and including population as a fixed effect. The ‘animal’-level random effect was distributed as *a N*(0, *G* ⊗ *A*) where *A* is the additive relatedness matrix derived from the breeding design and ⊗ is the Kronecker product, and yields the additive-genetic variance-covariance matrix *G* (Hadfield and Nakagawa, 2010). We sampled the posterior distribution for 75 000 iterations with a transient of 25 000 iterations discarded and a thinning interval of 50. We assessed and confirmed convergence by evaluating posterior trace plots and confirming effective sample sizes close to the expected 1000 posterior samples.

##### Measuring evolvability

Here, evolvability is defined as the mean-scaled additive genetic variance, giving the expected percent change in the trait mean per generation under a unit strength of selection (i.e. selection gradient; β) (Hansen *et al*., 2003). For multivariate phenotypes described by the additive-genetic variance matrix *G*, the mean evolvability *e* is given by the mean of the eigenvalues, which equals the mean of the univariate evolvabilities (Hansen and Houle, 2008). We also compared the multivariate autonomy (“independence”) of trait groups using R package “evolvability” (Hansen and Houle, 2008).

##### Analyses

Finally, we compared the estimated mean-scaled *P* and *G* in terms of their size (mean evolvability), and in terms of the expected magnitudes of the selection response they yield for a set of hypothetical selection gradients (“random skewers”) (Hansen and Houle, 2008). For this, we used 1000 random unit-length selection gradients. The latter approach compares the G-matrices in terms of the key parameters they are used to derive within the relevant theoretical framework (i.e. the response to selection). For each pairwise comparison of G-matrices, we calculated correlations over the posterior samples, and attained the median, mean and 95% confidence interval of the resulting correlation distribution.

## 5. RESULTS

### Variance and covariance of traits

We recorded trait values for floral traits flower corolla diameter, stigma length and anther length, as well as vegetative trait values for plant height, leaf length and leaf pair count. Recorded life-history and fitness traits included total flower count and flower phenology (timing of first flower). Species and population differences were most pronounced in vegetative traits and phenology (Figure 1).

**Figure 1.**
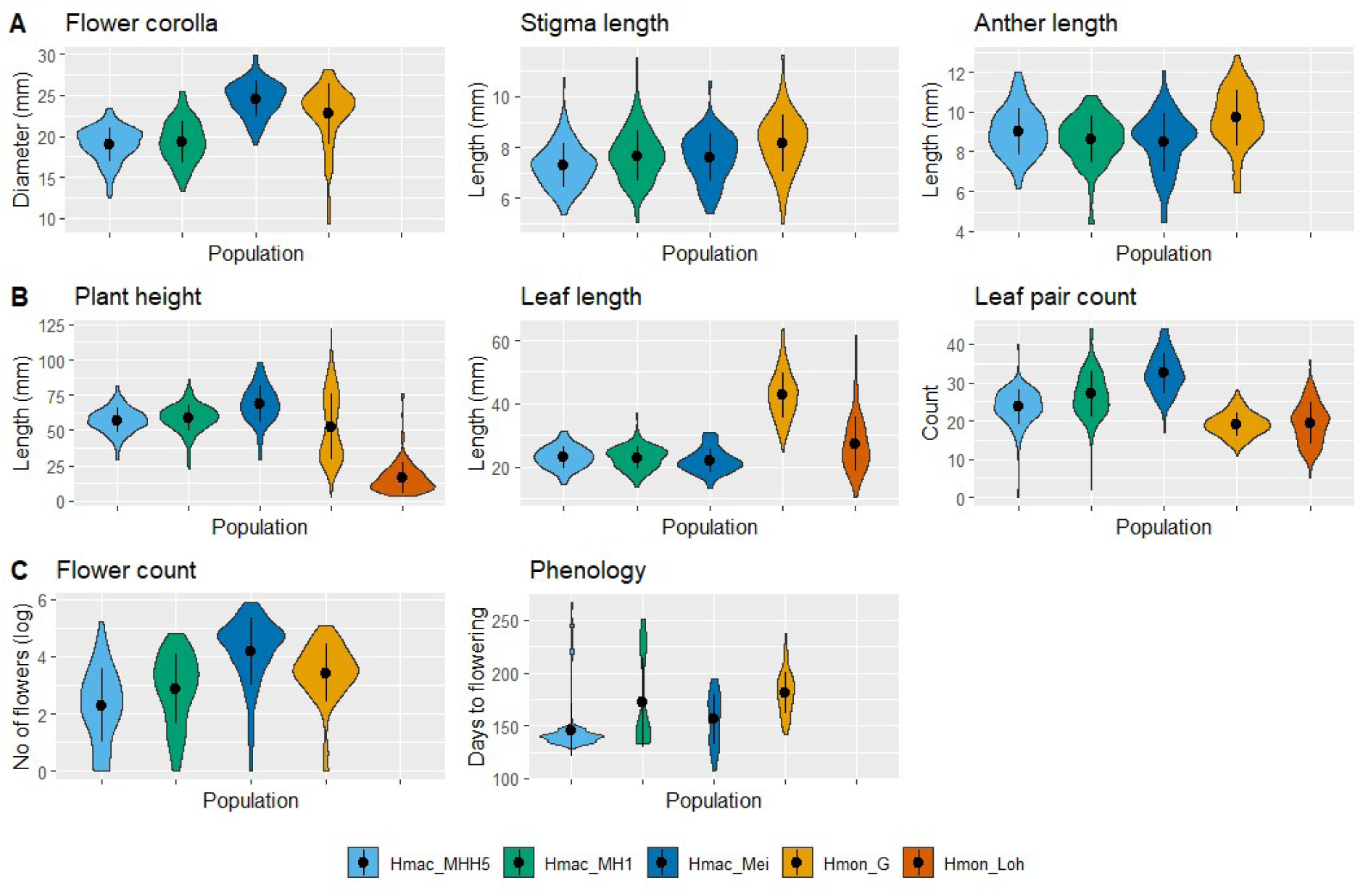
Distributions of measured traits in the five study populations. A) Floral traits, B) Vegetative traits, C) Life-history and fitness traits. For population Hmon_Loh, only vegetative trait data is available.

The mean-scaled phenotypic covariance matrices (*P*) of *H. maculatum* and *H. montanum* for vegetative, floral traits and life-history and fitness traits, using dataset 1, indicated that floral traits form a clear module of high covariance in both studied species (Table 2A). Furthermore, flower phenology showed positive covariance with leaf pair count and negative covariance with flower count in both species, and additionally negative covariance with plant height and leaf length in *H. montanum*. The *P* of vegetative traits using the larger dataset 2 revealed modularity within the recorded vegetative traits particularly in *H. montanum* (Table 2B). In *H. maculatum*, the strongest phenotypic covariance within vegetative traits was observed among plant height and leaf pair count, whereas leaf length was not strongly associated.

**Table 2.**
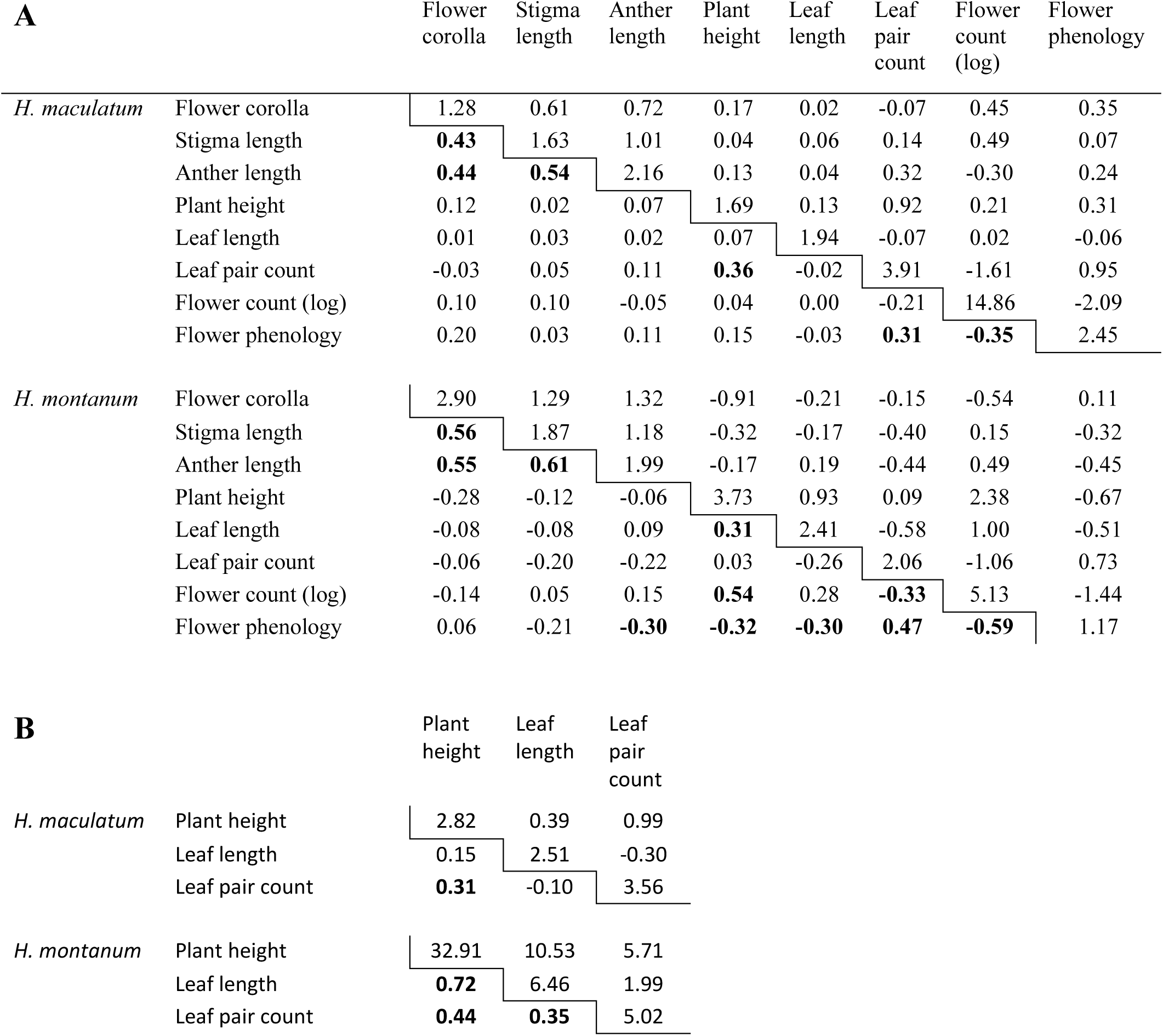
The mean-scaled phenotypic covariance matrices (*P*) of *H. maculatum* and *H. montanum* for A) vegetative, floral traits and life-history and fitness traits (Dataset 1) and B) only vegetative traits (Dataset 2), scaled by population trait mean and given as % (values on and above the diagonal). Values below the diagonal show the corresponding correlation matrix, with correlation values >0.3 shown in bold.

| A |  | Flower<br>corolla | Stigma<br>length | Anther<br>length | Plant<br>height | Leaf<br>length | Leaf<br>pair<br>count | Flower<br>count<br>(log) | Flower<br>phenology |
| --- | --- | --- | --- | --- | --- | --- | --- | --- | --- |
| *H. maculatum* | Flower corolla | 1.28 | 0.61 | 0.72 | 0.17 | 0.02 | -0.07 | 0.45 | 0.35 |
|  | Stigma length | **0.43** | 1.63 | 1.01 | 0.04 | 0.06 | 0.14 | 0.49 | 0.07 |
|  | Anther length | **0.44** | **0.54** | 2.16 | 0.13 | 0.04 | 0.32 | -0.30 | 0.24 |
|  | Plant height | 0.12 | 0.02 | 0.07 | 1.69 | 0.13 | 0.92 | 0.21 | 0.31 |
|  | Leaf length | 0.01 | 0.03 | 0.02 | 0.07 | 1.94 | -0.07 | 0.02 | -0.06 |
|  | Leaf pair count | -0.03 | 0.05 | 0.11 | **0.36** | -0.02 | 3.91 | -1.61 | 0.95 |
|  | Flower count (log) | 0.10 | 0.10 | -0.05 | 0.04 | 0.00 | -0.21 | 14.86 | -2.09 |
|  | Flower phenology | 0.20 | 0.03 | 0.11 | 0.15 | -0.03 | **0.31** | **-0.35** | 2.45 |
| *H. montanum* | Flower corolla | 2.90 | 1.29 | 1.32 | -0.91 | -0.21 | -0.15 | -0.54 | 0.11 |
|  | Stigma length | **0.56** | 1.87 | 1.18 | -0.32 | -0.17 | -0.40 | 0.15 | -0.32 |
|  | Anther length | **0.55** | **0.61** | 1.99 | -0.17 | 0.19 | -0.44 | 0.49 | -0.45 |
|  | Plant height | -0.28 | -0.12 | -0.06 | 3.73 | 0.93 | 0.09 | 2.38 | -0.67 |
|  | Leaf length | -0.08 | -0.08 | 0.09 | **0.31** | 2.41 | -0.58 | 1.00 | -0.51 |
|  | Leaf pair count | -0.06 | -0.20 | -0.22 | 0.03 | -0.26 | 2.06 | -1.06 | 0.73 |
|  | Flower count (log) | -0.14 | 0.05 | 0.15 | **0.54** | 0.28 | **-0.33** | 5.13 | -1.44 |
|  | Flower phenology | 0.06 | -0.21 | **-0.30** | **-0.32** | **-0.30** | **0.47** | **-0.59** | 1.17 |

| B |  | Plant<br>height | Leaf<br>length | Leaf<br>pair<br>count |
| --- | --- | --- | --- | --- |
| *H. maculatum* | Plant height | 2.82 | 0.39 | 0.99 |
|  | Leaf length | 0.15 | 2.51 | -0.30 |
|  | Leaf pair count | **0.31** | -0.10 | 3.56 |
| *H. montanum* | Plant height | 32.91 | 10.53 | 5.71 |
|  | Leaf length | **0.72** | 6.46 | 1.99 |
|  | Leaf pair count | **0.44** | **0.35** | 5.02 |

### Variation within and between species in univariate evolvability

Among populations, evolvability estimates varied to different extents depending on the G-matrix and included traits. The widest among-population variation in mean evolvability was found in ***G_LH_***, in which population mean evolvability varied between *e* = 1.23 - 3.06 (absolute difference in evolvability 1.83) (Table 6, Figure 2). G-matrix ***G_F+V_*** was most consistent across populations, with mean evolvability varying between e = 0.63 and e = 1.14 (absolute difference in evolvability 0.51). We found no consistent pattern for the magnitude of evolvability in the studied populations. For example, no particular population exhibited highest values of evolvability across the different G-matrices or traits. Furthermore, the two species showed on average relatively similar mean values of evolvability, with the exception of ***G_LH_***, in which mean evolvability was substantially higher in *H. maculatum* (e = 2.72) compared to *H. montanum* (e = 1.37) (99% higher, absolute difference in evolvability 1.35) (Table 6, Figure 2). Notably, evolvability generally varied more among populations within species than among the two different species. This was true also in ***G_V_***, estimated using our largest available dataset with multiple populations from both species. For example, the range of mean evolvability values of ***G_V_*** varied 0.66 units among populations of *H. montanum* and 0.73 units among populations of *H. maculatum*, whereas between species, mean evolvability varied only 0.14 units. The pattern of greater variation in evolvability among populations than among species was consistent also at the level of traits, i.e. in trait-specific evolvabilities.

**Figure 2.**
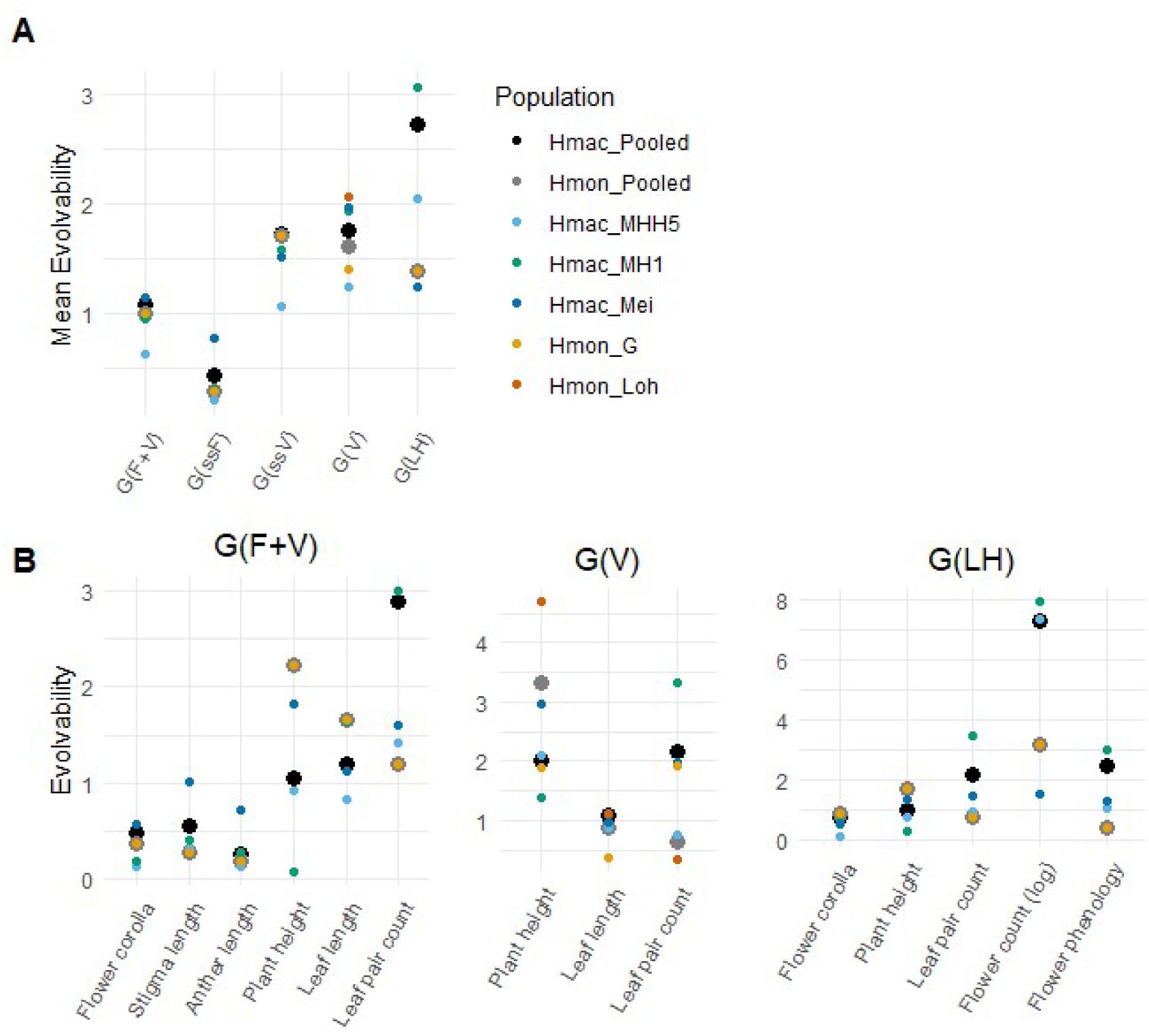
A) Mean evolvabilities of the different G-matrices for each species and population. B) Trait-specific evolvabilities for each species and population within the G-matrices ***G_F+V_***, ***G_V_*** and ***G_LH_***. Pooled data of study species *H. maculatum* is indicated by a large black dot and of *H. montanum* as a large gray dot. Population-specific data are indicated by small coloured dots. Data for population Hmon_Loh is available only for the estimation of G-matrix ***G_V_***.

### Evolvabilities and genetic covariances

The G-matrix ***G_F+V_*** including floral and vegetative traits (Table 3, Fig. 2) indicated substantially greater evolvability of vegetative than of floral traits in all study populations and in both studied *Hypericu*m species. In *H. maculatum* populations, the mean evolvability (*e*) of vegetative traits ranged from *e* = 1.06 to *e* = 1.58, and was 2-5 fold greater than that of floral traits (*e* = 0.20-0.77) (Table 6, Figure 2). In *H. montanum* (population Hmon_G), the evolvability of vegetative traits (*e* = 1.70) was 6-fold greater than the evolvability of floral traits (*e* = 0.28). In G-matrix ***G_V_*** using the largest available dataset and including only vegetative traits (Table 4), the mean evolvability estimates of vegetative traits were somewhat higher than in the same traits in ***G_F+V_*** estimated from a smaller dataset (including only flowering individuals), with *e* = 1.24-1.97 in *H. maculatum* populations (Table 6, Figure 2). Evolvability of vegetative traits in ***G_V_*** was similar also in *H. montanum* populations, with *e* = 1.39-2.05. Within vegetative traits, highest evolvability was found in *H. montanum* plant height (*e* = 3.31) (Table 4, Table 6, Figure 2). Overall, the highest mean evolvabilities were found for G-matrix ***G_LH_*** including life-history and fitness traits (Table 5), for which *e* varied between *e* = 1.24 and *e* = 3.06 in *H. maculatum* populations. In *H. montanum* (population Hmon_G), ***G_LH_*** had mean evolvability *e* = 1.37 (Table 6, Figure 2). For both species, particularly high values of mean evolvability were observed for flower count, with e = 7.29 in *H. maculatum* and e = 3.17 in *H. montanum* (Table 5, Figure 2). The variation in mean evolvabilities among populations correlated with the magnitude of evolvability. Thus, floral traits exhibited both the lowest values of evolvability and the least amounts of variation, whereas vegetative traits showed medium, and life-history traits the highest levels of evolvability and its among-population variability.

**Table 3.**
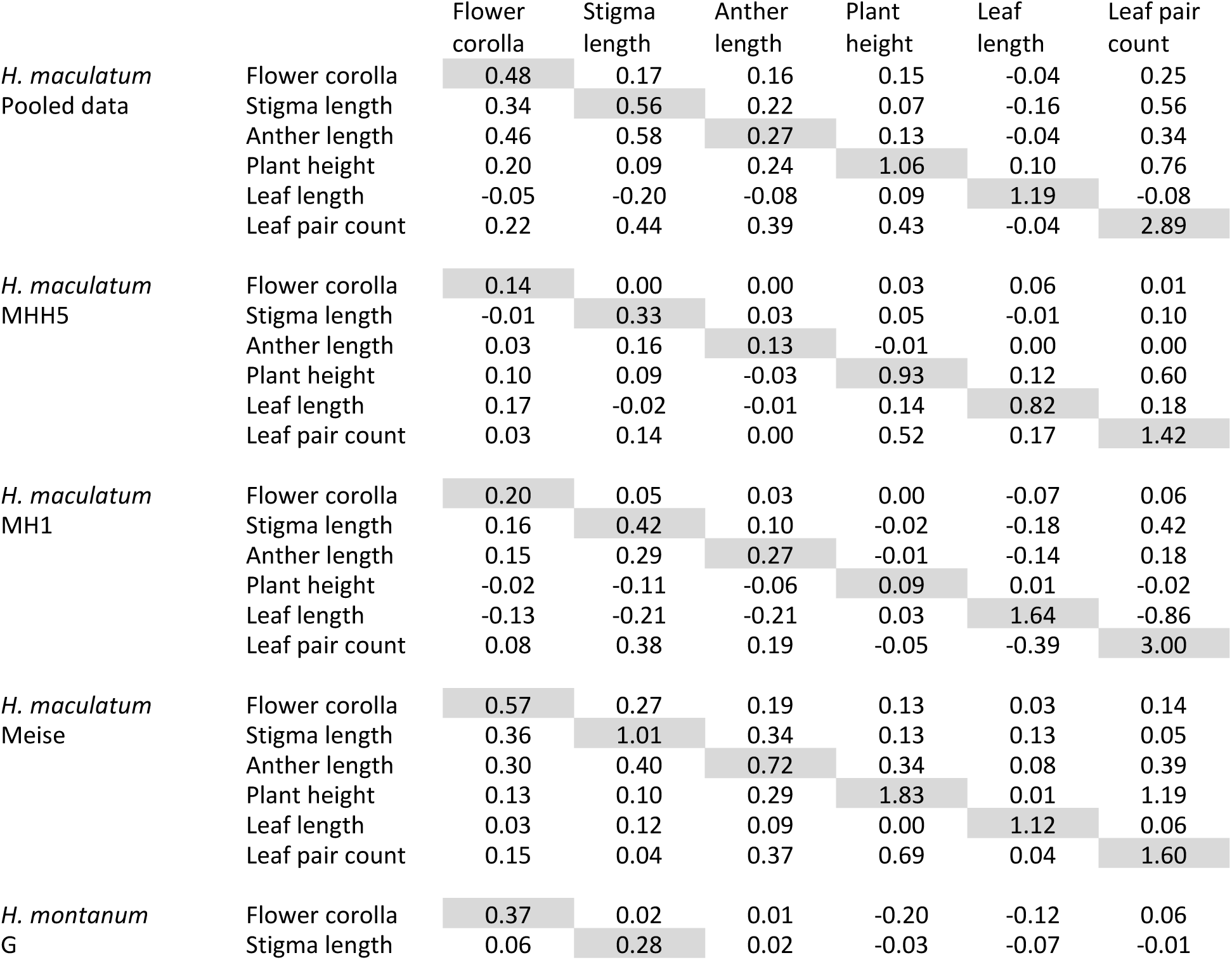
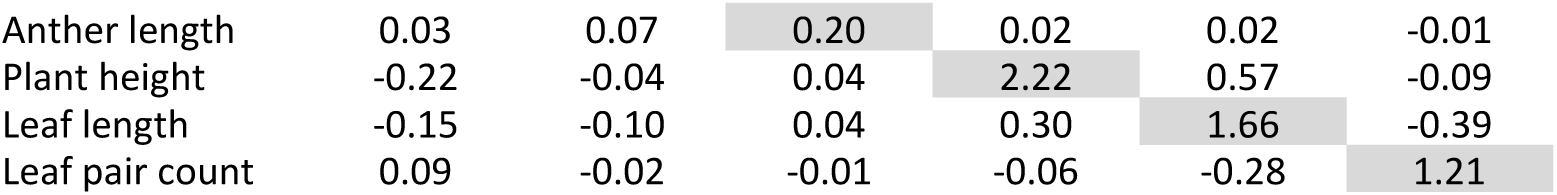
G-matrices of floral and vegetative traits (*G_F+V_*) in *H. maculatum* and *H. montanum* populations using dataset 1 (posterior median distribution). The upper triangle displays the genetic covariance matrix; the lower triangle displays the corresponding genetic correlation matrix. The shaded main diagonal reports trait-specific evolvability (mean-scaled genetic variance) for each trait.

|  |  | Flower<br>corolla | Stigma<br>length | Anther<br>length | Plant<br>height | Leaf<br>length | Leaf pair<br>count |
| --- | --- | --- | --- | --- | --- | --- | --- |
| *H. maculatum*<br>Pooled data | Flower corolla | 0.48 | 0.17 | 0.16 | 0.15 | -0.04 | 0.25 |
|  | Stigma length | 0.34 | 0.56 | 0.22 | 0.07 | -0.16 | 0.56 |
|  | Anther length | 0.46 | 0.58 | 0.27 | 0.13 | -0.04 | 0.34 |
|  | Plant height | 0.20 | 0.09 | 0.24 | 1.06 | 0.10 | 0.76 |
|  | Leaf length | -0.05 | -0.20 | -0.08 | 0.09 | 1.19 | -0.08 |
|  | Leaf pair count | 0.22 | 0.44 | 0.39 | 0.43 | -0.04 | 2.89 |
| *H. maculatum*<br>MHH5 | Flower corolla | 0.14 | 0.00 | 0.00 | 0.03 | 0.06 | 0.01 |
|  | Stigma length | -0.01 | 0.33 | 0.03 | 0.05 | -0.01 | 0.10 |
|  | Anther length | 0.03 | 0.16 | 0.13 | -0.01 | 0.00 | 0.00 |
|  | Plant height | 0.10 | 0.09 | -0.03 | 0.93 | 0.12 | 0.60 |
|  | Leaf length | 0.17 | -0.02 | -0.01 | 0.14 | 0.82 | 0.18 |
|  | Leaf pair count | 0.03 | 0.14 | 0.00 | 0.52 | 0.17 | 1.42 |
| *H. maculatum*<br>MH1 | Flower corolla | 0.20 | 0.05 | 0.03 | 0.00 | -0.07 | 0.06 |
|  | Stigma length | 0.16 | 0.42 | 0.10 | -0.02 | -0.18 | 0.42 |
|  | Anther length | 0.15 | 0.29 | 0.27 | -0.01 | -0.14 | 0.18 |
|  | Plant height | -0.02 | -0.11 | -0.06 | 0.09 | 0.01 | -0.02 |
|  | Leaf length | -0.13 | -0.21 | -0.21 | 0.03 | 1.64 | -0.86 |
|  | Leaf pair count | 0.08 | 0.38 | 0.19 | -0.05 | -0.39 | 3.00 |
| *H. maculatum*<br>Meise | Flower corolla | 0.57 | 0.27 | 0.19 | 0.13 | 0.03 | 0.14 |
|  | Stigma length | 0.36 | 1.01 | 0.34 | 0.13 | 0.13 | 0.05 |
|  | Anther length | 0.30 | 0.40 | 0.72 | 0.34 | 0.08 | 0.39 |
|  | Plant height | 0.13 | 0.10 | 0.29 | 1.83 | 0.01 | 1.19 |
|  | Leaf length | 0.03 | 0.12 | 0.09 | 0.00 | 1.12 | 0.06 |
|  | Leaf pair count | 0.15 | 0.04 | 0.37 | 0.69 | 0.04 | 1.60 |
| *H. montanum*<br>G | Flower corolla | 0.37 | 0.02 | 0.01 | -0.20 | -0.12 | 0.06 |
|  | Stigma length | 0.06 | 0.28 | 0.02 | -0.03 | -0.07 | -0.01 |

|  |  |  |  |  |  |  |
| --- | --- | --- | --- | --- | --- | --- |
| Anther length | 0.03 | 0.07 | 0.20 | 0.02 | 0.02 | -0.01 |
| Plant height | -0.22 | -0.04 | 0.04 | 2.22 | 0.57 | -0.09 |
| Leaf length | -0.15 | -0.10 | 0.04 | 0.30 | 1.66 | -0.39 |
| Leaf pair count | 0.09 | -0.02 | -0.01 | -0.06 | -0.28 | 1.21 |

**Table 4.** G-matrices of vegetative traits (*G_V_*) in the largest available dataset (dataset 2) in *H. maculatum* and *H. montanum* populations (posterior median distribution, flowering (y/n) included as a fixed effect). The upper triangle displays the genetic covariance matrix; the lower triangle displays the corresponding genetic correlation matrix. The shaded main diagonal reports trait-specific evolvability (mean-scaled genetic variance) for each trait.

|  |  | Plant height | Leaf length | Leaf pair count |
| --- | --- | --- | --- | --- |
| *H. maculatum*<br>Pooled data | Plant height | 2.02 | 0.10 | 1.10 |
|  | Leaf length | 0.07 | 1.08 | -0.43 |
|  | Leaf pair count | 0.53 | -0.28 | 2.16 |
| *H. montanum*<br>Pooled data | Plant height | 3.31 | 0.99 | 0.90 |
|  | Leaf length | 0.58 | 0.88 | 0.18 |
|  | Leaf pair count | 0.62 | 0.25 | 0.64 |
| *H. maculatum*<br>MHH5 | Plant height | 2.09 | 0.44 | 0.88 |
|  | Leaf length | 0.32 | 0.87 | 0.08 |
|  | Leaf pair count | 0.70 | 0.10 | 0.75 |
| *H. maculatum*<br>MH1 | Plant height | 1.37 | -0.16 | 0.58 |
|  | Leaf length | -0.13 | 1.10 | -0.90 |
|  | Leaf pair count | 0.27 | -0.47 | 3.33 |
| *H. maculatum*<br>Mei | Plant height | 2.97 | -0.05 | 1.85 |
|  | Leaf length | -0.03 | 0.97 | 0.00 |
|  | Leaf pair count | 0.77 | 0.00 | 1.97 |
| *H. montanum*<br>G | Plant height | 1.88 | 0.20 | 0.08 |
|  | Leaf length | 0.24 | 0.37 | -0.12 |
|  | Leaf pair count | 0.04 | -0.15 | 1.91 |
| *H. montanum*<br>Loh | Plant height | 4.69 | 0.81 | 0.54 |
|  | Leaf length | 0.35 | 1.13 | 0.09 |
|  | Leaf pair count | 0.43 | 0.15 | 0.34 |

**Table 5.** G-matrices of life-history and fitness traits (*G_LH_*) in *H. maculatum* and *H. montanum* populations using dataset 1 (posterior median distribution). The upper triangle displays the genetic covariance matrix; the lower triangle displays the corresponding genetic correlation matrix. The shaded main diagonal reports trait-specific evolvability (mean-scaled genetic variance) for each trait.

|  |  | Flower corolla | Plant height | Numb leaf pairs | Flower count (log) | Flower phenology |
| --- | --- | --- | --- | --- | --- | --- |
| *H. maculatum* | Flower corolla | 0.73 | 0.08 | 0.22 | -0.12 | 0.35 |
| Pooled data | Plant height | 0.10 | 0.96 | 0.67 | -0.89 | 0.15 |
|  | Numb leaf pairs | 0.18 | 0.46 | 2.16 | -1.59 | 0.88 |
|  | Flower count (log) | -0.05 | -0.33 | -0.40 | 7.29 | -2.19 |
|  | Flower phenology | 0.26 | 0.10 | 0.38 | -0.52 | 2.46 |
| *H. maculatum*<br>MHH5 | Flower corolla | 0.13 | 0.03 | 0.01 | 0.11 | -0.02 |
|  | Plant height | 0.09 | 0.78 | 0.42 | -0.20 | -0.02 |
|  | Numb leaf pairs | 0.03 | 0.50 | 0.90 | -0.06 | -0.13 |
|  | Flower count (log) | 0.11 | -0.08 | -0.02 | 7.35 | -1.17 |
|  | Flower phenology | -0.06 | -0.02 | -0.13 | -0.42 | 1.05 |
| *H. maculatum*<br>MH1 | Flower corolla | 0.57 | -0.01 | 0.17 | -0.48 | 0.50 |
|  | Plant height | -0.02 | 0.27 | 0.00 | -0.13 | -0.05 |
|  | Numb leaf pairs | 0.12 | 0.00 | 3.48 | -1.83 | 1.71 |
|  | Flower count (log) | -0.23 | -0.09 | -0.35 | 7.97 | -2.67 |
|  | Flower phenology | 0.38 | -0.05 | 0.53 | -0.55 | 2.99 |
| *H. maculatum*<br>Mei | Flower corolla | 0.55 | 0.21 | 0.22 | 0.00 | 0.13 |
|  | Plant height | 0.24 | 1.37 | 1.01 | -0.28 | 0.61 |
|  | Numb leaf pairs | 0.24 | 0.71 | 1.47 | -0.36 | 0.69 |
|  | Flower count (log) | 0.00 | -0.19 | -0.24 | 1.51 | -0.42 |
|  | Flower phenology | 0.16 | 0.46 | 0.50 | -0.30 | 1.29 |
| *H. montanum*<br>G | Flower corolla | 0.88 | -0.34 | 0.06 | -0.29 | 0.09 |
|  | Plant height | -0.28 | 1.70 | -0.14 | 1.24 | -0.33 |
|  | Numb leaf pairs | 0.07 | -0.13 | 0.73 | -0.59 | 0.20 |
|  | Flower count (log) | -0.18 | 0.53 | -0.39 | 3.17 | -0.58 |
|  | Flower phenology | 0.16 | -0.40 | 0.38 | -0.52 | 0.39 |

**Table 6:** Population-specific evolvability values (e_mean (shaded): average unconditional evolvability, e_min: minimum evolvability, e_max: maximum evolvability, r_mean: average respondability, c_mean: average conditional evolvability). and estimates of multivariate autonomy (a_mean: average autonomy, i_mean: average integration) of the studied G-matrices, as defined in Hansen and Houle (2008).

| Population | G-matrix | e_mean | e_min | e_max | c_mean | r_mean | a_mean | i_mean |
| --- | --- | --- | --- | --- | --- | --- | --- | --- |
| Hmac Pooled | $G_{F+V}$ | 1.07 | 0.13 | 3.37 | 0.51 | 1.36 | 0.57 | 0.43 |
| | $G_{ssF}$ | 0.43 | 0.13 | 0.83 | 0.33 | 0.48 | 0.87 | 0.13 |
| | $G_{ssV}$ | 1.71 | 0.75 | 3.16 | 1.40 | 1.84 | 0.92 | 0.08 |
| Hmac MHH5 | $G_{F+V}$ | 0.63 | 0.12 | 1.87 | 0.32 | 0.78 | 0.58 | 0.42 |
| | $G_{ssF}$ | 0.20 | 0.12 | 0.34 | 0.18 | 0.21 | 0.95 | 0.05 |
| | $G_{ssV}$ | 1.06 | 0.53 | 1.87 | 0.91 | 1.12 | 0.94 | 0.06 |
| Hmac MH1 | $G_{F+V}$ | 0.94 | 0.08 | 3.51 | 0.29 | 1.32 | 0.43 | 0.57 |
| | $G_{ssF}$ | 0.30 | 0.18 | 0.48 | 0.27 | 0.31 | 0.96 | 0.04 |
| | $G_{ssV}$ | 1.58 | 0.09 | 3.42 | 0.45 | 1.88 | 0.36 | 0.64 |
| Hmac Mei | $G_{F+V}$ | 1.14 | 0.39 | 3.07 | 0.76 | 1.34 | 0.74 | 0.26 |
| | $G_{ssF}$ | 0.77 | 0.43 | 1.37 | 0.65 | 0.81 | 0.94 | 0.06 |
| | $G_{ssV}$ | 1.52 | 0.52 | 2.91 | 1.17 | 1.66 | 0.88 | 0.12 |
| Hmon G | $G_{F+V}$ | 0.99 | 0.19 | 2.67 | 0.51 | 1.22 | 0.58 | 0.42 |
| | $G_{ssF}$ | 0.28 | 0.19 | 0.37 | 0.27 | 0.29 | 0.99 | 0.01 |
| | $G_{ssV}$ | 1.70 | 0.94 | 2.64 | 1.56 | 1.76 | 0.96 | 0.04 |
| Hmac Pooled | $G_V$ | 1.75 | 0.66 | 3.22 | 1.34 | 1.95 | 0.79 | 0.21 |
| Hmon Pooled | $G_V$ | 1.61 | 0.31 | 3.92 | 0.70 | 2.09 | 0.52 | 0.48 |
| Hmac MHH5 | $G_V$ | 1.24 | 0.29 | 2.63 | 0.72 | 1.48 | 0.64 | 0.36 |
| Hmac MH1 | $G_V$ | 1.93 | 0.78 | 3.79 | 1.41 | 2.19 | 0.78 | 0.22 |
| Hmac Mei | $G_V$ | 1.97 | 0.55 | 4.38 | 1.13 | 2.40 | 0.65 | 0.35 |
| Hmon G | $G_V$ | 1.39 | 0.33 | 1.98 | 0.96 | 1.54 | 0.68 | 0.32 |
| Hmon Loh | $G_V$ | 2.05 | 0.27 | 4.93 | 0.81 | 2.65 | 0.47 | 0.53 |
| Hmac Pooled | $G_{LH}$ | 2.72 | 0.56 | 8.78 | 1.30 | 3.52 | 0.63 | 0.37 |
| Hmac MHH5 | $G_{LH}$ | 2.04 | 0.13 | 7.57 | 0.59 | 2.91 | 0.43 | 0.57 |
| Hmac MH1 | $G_{LH}$ | 3.06 | 0.27 | 10.04 | 0.94 | 4.11 | 0.41 | 0.59 |
| Hmac Mei | $G_{LH}$ | 1.24 | 0.41 | 3.20 | 0.81 | 1.45 | 0.75 | 0.25 |
| Hmon G | $G_{LH}$ | 1.37 | 0.24 | 4.17 | 0.74 | 1.72 | 0.67 | 0.33 |

Among the measured floral traits, genetic covariance was indicated for some populations and pooled data in *H. maculatum*, but not for *H. montanum* (Table 3). Genetic covariance among the different vegetative traits was also observed, particularly involving leaf pair count, which showed positive genetic covariance with plant height in both species and additionally negative covariance with leaf length in *H. maculatum* (Table 4). In *H. maculatum*, leaf pair count also had positive genetic covariance with floral measures (Table 3). ***G_LH_*** revealed negative genetic covariance between flower count and leaf pair count for all studied populations (Table 5). Flower count also had negative genetic covariance with plant height in *H. maculatum* (in which the two vegetative traits leaf pair count and plant height had positive genetic covariance), but positive genetic covariance with plant height in *H. montanum* (population Hmon_G). Finally, flower phenology had substantial negative covariance with flower count in all studied populations (Table 5).

### Similarity of G-matrices among species and populations

Pairwise correlations of G-matrices showed varying levels of similarity in genetic architecture within and among species (Figure 3, Table S2). Correlation values were substantially greater when comparing posterior distribution medians and means, compared to comparisons of posterior pairwise samples (Table S2), which could reflect high uncertainty in the estimates. This is also indicated by the wide confidence intervals of pairwise sample correlations, which consistently cross zero (Figure 3). Here, we focus on the mean correlations over posterior samples.

**Figure 3.**
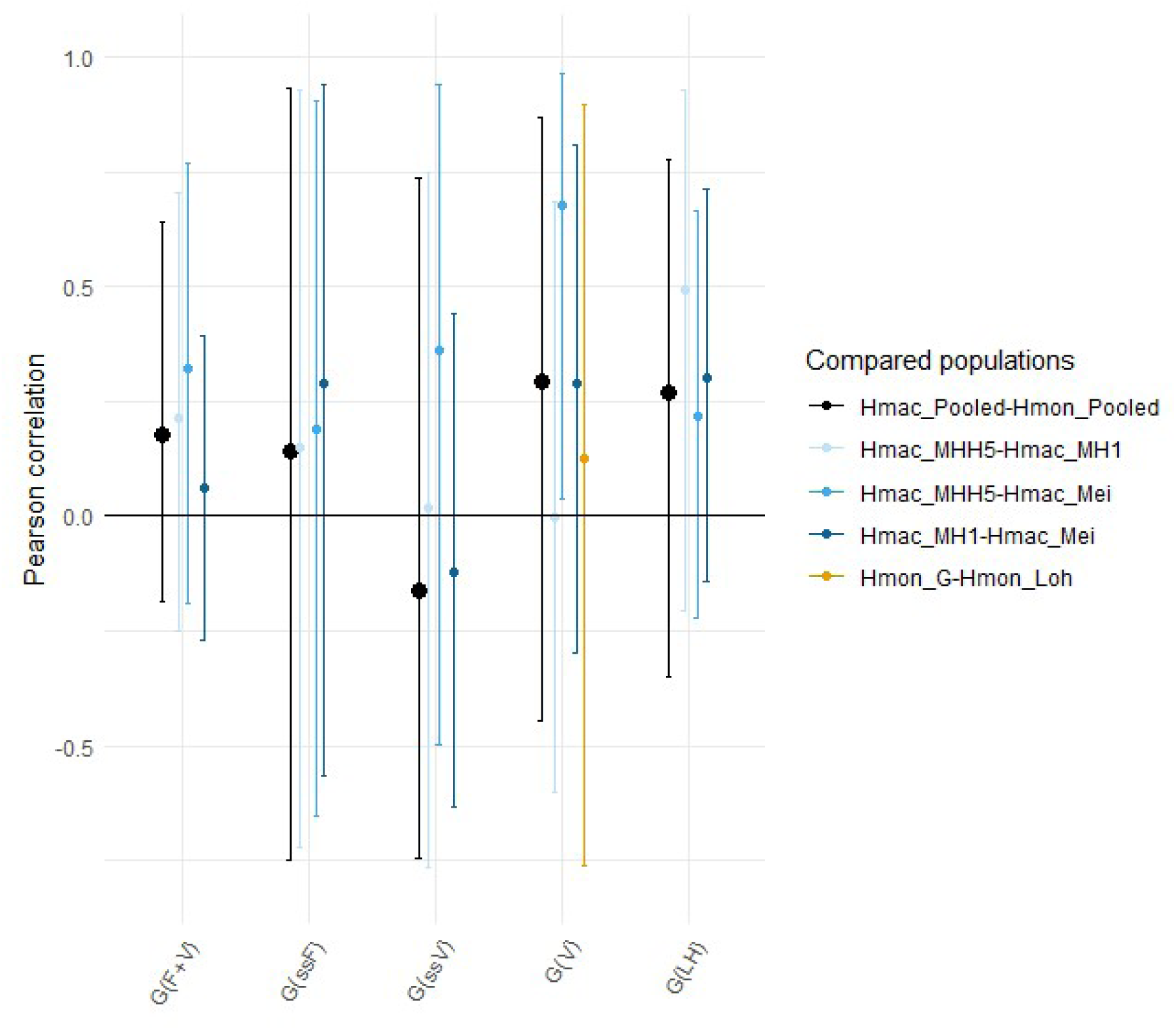
Pairwise comparisons of G-matrices ***G_F+V_*** (floral and vegetative traits), ***G_ssF_*** (floral traits subset) ***G_ssV_*** (vegetative traits subset) ***G_V_*** (vegetative traits) and ***G_LH_*** (life-history traits) through 1000 simulated values of evolvability along random selection gradients given the derived G-matrices of *H. maculatum* and *H. montanum* study populations. The correlations are calculated over pairwise posterior samples. Dots represent mean Pearson correlations, and whiskers the 95% confidence intervals. Interspecific comparisons of G-matrices, pooling data per species, are indicated by large black dots and black whiskers.

In general, the similarity of *H. maculatum* G-matrices varied between cor = -0.12 – 0.68 depending on the G-matrix traits and compared populations (Figure 3). Notably, there was no consistent pattern of particular population pairs having the most similar expected evolutionary trajectories, but similarity varied between trait classes. Among the trait classes, life history traits showed on average highest similarity in G-matrices among the three different *H. maculatum* populations, with average cor = 0.34 in pairwise samples among population-specific ***G_LH_*** (Figure 3). ***G_LH_*** showed also the highest correlation between the two species *H. maculatum* and *H. montanum* (interspecific cor = 0.27), together with ***G_V_*** comprised of vegetative traits (interspecific cor = 0.29).

Floral traits seemed to show higher intra- and interspecific similarity in genetic architecture than did vegetative traits, with ***G_ssF_*** average cor = 0.21 in pairwise samples among *H. maculatum* populations, and interspecific cor = 0.14. This contrasts with the lower correlations of ***G_ssV_*** (vegetative trait subset for flowering individuals) among *H. maculatum* populations (average cor = 0.09), and even a negative correlation among the two species (cor = - 0.16). However, we did not find a respective negative interspecific correlation for ***G_V_*** comprising vegetative traits in the most extensive dataset including both flowering and non-flowering individuals, for which interspecific cor = 0.29. Among *H. maculatum* populations, the correlation of vegetative trait genetic architecture (***G_V_***) varied particularly widely (cor = -0.00 - 0.68). Furthermore, the correlation of ***G_V_*** among the two *H. montanum* populations (cor = 0.13) was lower than the correlation of ***G_V_*** among the two species (cor = 0.29). A lower intraspecific correlation compared to the respective interspecific one was found also in some other population pairwise comparisons (Figure 3). However, on average, intraspecific correlations of G-matrices (averag cor = 0.22) were higher than interspecific correlations (average cor = 0.14).

### Correlations of G- and P-matrices

Overall, G-matrices were highly positively correlated with their respective P-matrices, with cor = 0.57 – 0.96 (Figure 4, Table S3). The highest correlation of *G and P* was found for ***G_LH_***, composed of life-history traits, for which the average correlation of respective *G and P* reached a high value of cor = 0.96. For vegetative traits, correlation of *G and P* was also high (cor = 0.93 and cor = 0.79 for G-matrices ***G_ssV_*** and ***G_V_***, respectively; Table S3, Figure 4). The correlation of *G and P*was lowest for floral traits, for which average cor = 0.57 (***G_ssF_***).

**Figure 4.**
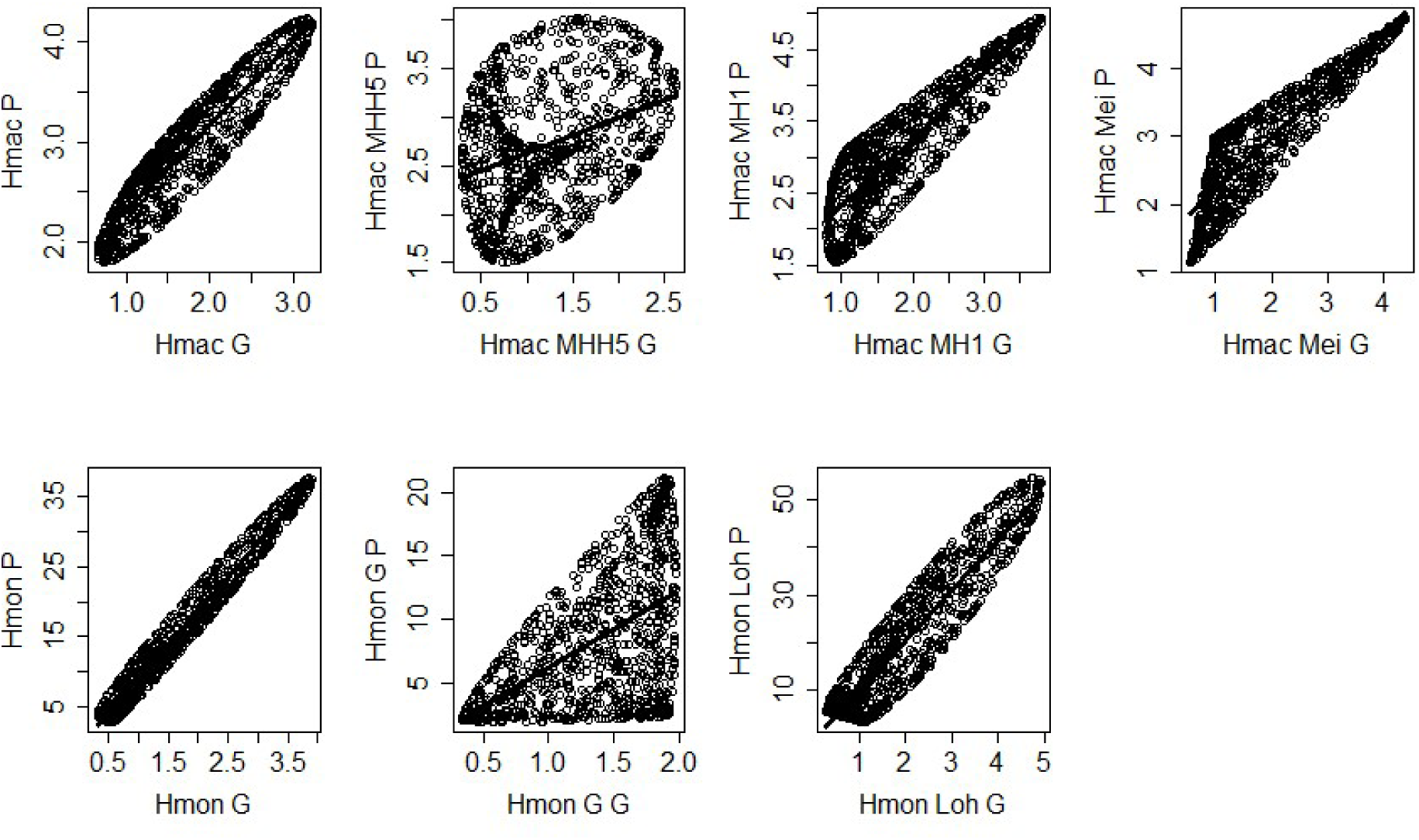
Pairwise comparisons of vegetative trait G-matrices ***G_F+V_*** and the respective P-matrices through 1000 simulated values of evolvability along random selection gradients given the derived G- and respective P-matrices of studied *H. maculatum* and *H. montanum* populations. The upper panel shows comparisons for *H. maculatum*, and lower panel for *H. montanum*. The leftmost plots show species-specific comparisons of G- and P-matrices, pooling data per species.

## 6. DISCUSSION

One of the most pressing challenges for predicting the consequences of global change on biodiversity is the need for empirical measures of additive genetic variances, evolvabilities and genetic variance-covariance matrices (i.e. G-matrices) for more species, populations, and traits (Hoffmann and Sgrò, 2011; Urban *et al*., 2023). Here, we provide such data for multiple populations of two congeneric species of one of the most understudied groups of organisms in this regard, i.e. vascular plants. We discuss variation in evolvability across populations and species, patterns of evolvability across traits, variation in G-matrix structure, as well as scaling between genetic and phenotypic variances.

### Variation of evolvability within and between species

Overall levels of evolvability in the two studied *Hypericum* species were somewhat lower than found on average in previous studies on plants (median evolvabilities of floral and vegetative traits in a meta-analysis of 54 plant taxa e = 0.98% and e = 2.08%, respectively (Opedal, 2019), compared with e = 0.36% and e = 1.68% measured here). These values indicate relatively limited evolutionary potential. For example, a 50 % change in a floral or vegetative trait would require on average 140 or 30 generations of evolution under consistent strong selection (β = 1), respectively.

On the other hand, we found substantial variation in mean evolvability among populations. Interestingly, evolvabilities were more variable among populations within species than among the two different species. Furthermore, we found no clear pattern for more similar evolvabilities in e.g. populations with closer geographic origin, or consistent relative magnitude of evolvability in populations across the different G-matrices. That is, populations with the highest evolvabilities varied depending on the G-matrix. The overall interpretation of this pattern highlights the dynamic nature of evolutionary processes at different biological scales. It suggests that within-species adaptation to local environments can drive significant variational changes in evolvability, possibly more so than seen at the species level where broader evolutionary constraints may play a more homogenizing role (Langerhans and DeWitt, 2004; Hereford, 2009). Species may share similar functional or developmental constraints due to evolutionary history that can limit differences in evolvability compared to the more varied scenarios encountered by populations within a species.

For the potential of climate adaptation, this implies that due to varying adaptive capacity of populations within a species, some populations may have greater chances of evolutionary rescue with changing conditions. Furthermore, it highlights the importance of population choice in conservation actions such as translocations. However, with seemingly stochastic variation in evolvability among populations, it remains a challenge to predict evolvability in a particular population without direct empirical data.

### Patterns of evolvability across traits

We found that both the magnitude and variability of evolvability estimates varied among G-matrices and traits. Floral traits exhibited both the lowest values of evolvability, as well as the least amounts of variation, whereas vegetative traits showed medium, and life-history traits the highest levels of evolvability and its variability. The comparatively low evolvability of floral traits, particularly those closely associated with pollination mechanisms, is emerging as a consistent trend across plant species (Opedal, 2019). It is thought to reflect historical pollinator-mediated stabilizing selection, thus highlighting the role of pollinators in the evolution of floral traits. Although evolvability of floral traits may allow evolutionary responses to often strong selection pressures imposed by pollinator interactions, the lower evolutionary potential of flowers compared to other trait types may to some extent become a key limiting factor for the adaptation of plants to global change (Mattila *et al*., 2024), particularly to climate-driven shifts in pollinator communities and global pollinator decline (Potts *et al*., 2010).

The high levels of evolvability and its variability in life-history traits could be explained by their direct or close connection with fitness. Fitness traits often have higher levels of additive genetic variance than traits not closely linked with fitness (Merilä and Sheldon, 1999). This is supported by our observation of the highest values of evolvability for *Hypericum* flower abundance, a close proxy of reproductive output. The genetic architecture underpinning fitness traits can be complex, which can both promote high levels of evolvability and lead to variation in evolvability among populations under differential selection pressures (Merilä and Sheldon, 1999; Roff and Fairbairn, 2007). Furthermore, given their strong connection to reproductive success and thus potential for strong selection gradients, fitness traits can evolve relatively quickly in response to changing conditions and differing ecological pressures, contributing to greater among-population variation.

In contrast, morphological traits, which are often under weak optimizing selection, tend to have lower evolvability (Houle, 1992; Hansen, Pélabon and Houle, 2011). According to Hansen et al. (2011), the median evolvability for life-history traits is about 1%, while for linear size traits it is about 0.1%.

Comparative values observed for *Hypericum* populations were somewhat higher, nonetheless demonstrating significantly lower evolvability for vegetative size traits (*H. maculatum* e = 1.75%) than life-history traits (*H. maculatum* e = 2.72%). The selection pressures on morphological traits may consist of many different types of selection, particularly those driven by abiotic environmental conditions, but also include selection through predation/herbivory, competition and resource availability (Kingsolver *et al*., 2001; Flatt and Heyland, 2011). In plants, vegetative traits such as leaf size are often largely driven by local environmental conditions, e.g., temperature, light availability and moisture (Caruso, Maherali and Martin, 2020). This could lead to moderate variation in evolvability among populations that have adapted to differing environmental conditions. Furthermore, vegetative traits often display phenotypic plasticity, allowing them to adapt to their immediate environment even with limited genetic variation. This plasticity might result in moderate genetic differences among populations as they adapt over time.

### Variation in G-matrix structure

The stability of **G** between species and among populations of a species remains a topic of ongoing research. For example, while it’s been suggested that populations at the species’ range edges could differ in genetic diversity and evolutionary potential compared to those in the core areas, empirical evidence remains limited (Eckert, Samis and Lougheed, 2008). Some studies suggest that the G-matrix is capable of rapid evolution across generations and populations (see e.g. (Björklund, Husby and Gustafsson, 2013). However, the field is challenged by the use of diverse methods for comparing G-matrices, particularly analysing variance-scaled data, which can skew or blur effects (Hansen, Pélabon and Houle, 2011). Overall, particularly for morphological traits, it has been generally assumed that G-matrices are relatively stable within species (Puentes, Granath and Ågren, 2016; McGlothlin *et al*., 2022; Henry and Stinchcombe, 2023; Mattila *et al*., 2024). A major aim of the current study is to provide comparable evidence to help resolve this largely uncharted question.

Against our expectations, the pairwise correlations between G-matrices, as estimated through simulated responses to random selection gradients, indicated only moderate similarity of **G** within and among species (mean cor = -0.12 – 0.68 depending on the G-matrix traits and compared populations). However, these correlation values come with wide error margins, demonstrating the difficulty of obtaining reliable empirical estimates of **G**. Intraspecific correlations (mean cor = 0.22) were somewhat higher than interspecific ones (mean cor = 0.14), but both the level of these correlations and the difference between intra- and interspecific correlations was less than expected. We also did not find any consistent patterns of particular population pairs having the most similar evolutionary trajectories, but similarity varied between trait classes. Together, these results suggest that genetic architecture may be quite labile across species and populations. Across species distributions, differentially correlating selection pressures in different trait types may result in various correlational patterns in genetic architecture across the range.

We expected to find particularly conserved genetic architecture in floral traits, which have often evolved under strong stabilizing selection and can exhibit strong patterns of covariance among traits (Berg, 1960; Opedal, 2019). When studying only flowering individuals, the results indeed indicated higher intra- and interspecific similarity in floral than vegetative trait genetic architecture. However, vegetative trait G-matrix similarity was higher when including also non-flowering individuals.

Furthermore, the highly variable pairwise similarity of vegetative trait ***G_V_*** was surprising, although we also found some high ***G_V_*** intraspecific pairwise correlation values (up to cor = 0.68), indicating that some conspecific populations share highly similar genetic architecture underlying these traits. On average, the highest similarity in G-matrix structure among conspecific populations was found for life-history traits. This suggests relatively correlated evolutionary trajectories of this trait group among populations, despite the expected complex genetic underpinnings (Merilä and Sheldon, 1999; Roff and Fairbairn, 2007) and observed high variability in population mean evolvability of life-history trait ***G_LH_***.

### Scaling between genetic and phenotypic variances

The estimation of G-matrices using empirical data with multigenerational pedigree information is notoriously tedious, and for many taxa, unfeasible. However, recent important advances in the field of quantitative genetics indicate that direct genetic data may be unnecessary for estimating evolutionary potential in many cases (Holstad *et al*., 2024). This is because phenotypic and genetic variation have been found to commonly scale near-isometrically, suggesting that G-matrices can often be inferred directly from phenotypic variance matrices (i.e. P-matrices), with no need for associated pedigree data. Our results support this observation, with overall high correlations found between estimated G-matrices and respective P-matrices, commonly approaching isometric scaling (cor = 1). On the other hand, although mean correlations were high, there was also significant variation in the scaling between ***G*** and ***P*** among populations and among G-matrices, with some correlation values as low as around cor = 0.6. This suggests that while genetic factors largely mirror phenotypic variation, discrepancies may arise, potentially due to e.g. genotype-by-environment interactions, epigenetic modifications and non-additive genetic effects (dominance and epistasis). Overall, we found the weakest scaling between ***G*** and ***P*** in floral traits, which may reflect the overall low additive genetic variance in these traits.

### Can evolutionary potential help plants survive climate change?

Because evolvability can be translated into expected trait change per generation under specified selection, it offers a practical framework for predicting how species might evolve in response to global change (Urban *et al*., 2023; Mattila *et al*., 2024). Recent research demonstrating strong positive correlations between microevolutionary evolvability and long-term population divergence underscores its predictive value (Opedal *et al*., 2023; Holstad *et al*., 2024). The applicability of evolvability estimates for evaluating the adaptive potential in threatened populations was demonstrated in a recent study on the arctic Siberian primrose (*Primula nutans*) (Mattila *et al*., 2024). The study indicated that the rate of environmental change is likely to by far outpace the rate of evolution plausible for the species. In the light of the above results, it is notable that the evolvability values estimated here for the more commonly occurring and widely-distributed *Hypericum* species were, in most traits, similar in magnitude compared with those in the endangered *P. nutans* (vegetative trait *e* = 2.5 *vs. e* = 1.8, floral trait *e* = 0.2 *vs. e* = 0.4, and life-history trait *e* = 0.4 *vs. e* = 2.5 in *P. nutans* and *H. maculatum*, respectively). On the other hand, also substantially higher evolvabilities were observed for *Hypericum* (particularly in life-history traits), which may provide important adaptive opportunities. Consistent with *P. nutans* and previous studies on plants (Opedal, 2019), evolvability in *Hypericum* is most restricted in floral traits, which could become a limiting factor for plant responses to climate-driven threats, particularly shifts in pollinator communities.

However, unlike the primrose, which is geographically strictly confined to Fennoscandian seashores, the studied *Hypericum* may be able to respond to changing climate also by tracking suitable conditions with distributional shifts. Given the accumulating evidence indicating generally limited potential for evolution in natural populations to track rapidly shifting adaptive optima driven by climate change (Diniz-Filho and Bini, 2019; Morgan *et al*., 2020; Mattila *et al*., 2024), it will be crucial to maintain the capacity for distributional shifts e.g. by conserving natural habitat (Lawlor *et al*., 2024).

However, the main objective of our current study was not to evaluate climate-adaptive potential in *Hypericum* per se., but to evaluate overall intra- and interspecific variability in evolvability and G-matrix structure, which was observed to be unexpectedly high. This observation indicates that predicting population genetic variances based on e.g. taxonomic relationship, population size or geographic location may generally be difficult, and that reliable estimates require relevant empirical data.

Despite being highly interpretable, quantitative-genetic estimates of evolvability have been infrequently employed in studies examining adaptation to environmental changes, owing largely to the deficiency in available empirical estimates (Urban *et al*., 2023). Our study takes steps to bridge this gap, both in providing comparable estimates for predictive studies, in assessing the variability in evolvability and G-matrix structures within and among species, as well as motivating the acquisition of empirical genetic variance estimates for an increasing number of taxa. The promise of a potential “shortcut” to G-matrix estimates using straightforward phenotypic trait data calls for further investigation, and may provide an avenue for building a much needed library of data on evolutionary potentials in natural populations. A further challenge will be to connect genetic variance and covariance estimates to climatic selection pressures and climate gradients to enable predictions on the responses of biodiversity to ongoing climate change (Urban *et al*., 2023).

## ACKNOWLEDGEMENTS

Mari Miranto, Marija Bučar, Favour Mba, Shiromi Samiraja, Martti Levo, Laura Koivuniemi and Julia Lemmetty are thanked for technical assistance in the experiments. Sandrine Godefroid collected seeds of *H. maculatum* in Belgium. Tarja Niemelä provided plastic pollination bags. AM, LP, CM and MHH were funded by the Research Council of Finland (grants 331527 and 330739), and AM also by the Kone Foundation. ØO was supported by the Swedish Research Council (grant nr. 2021-04777) and the Crafoord Foundation (grant nr. 20210661), MTH by the Finnish Museum of Natural History, SK by Societas pro Fauna et Flora Fennica, Waldemar von Frenckells Stiftelse, Nordenskiöld-Samfundet and Kone Foundation, LP by the Jenny and Antti Wihuri Foundation, and MHH by the Jane and Aatos Erkko Foundation through the Research Centre for Ecological Change, University of Helsinki. MHH also acknowledges funding from the Research Council of Finland (grant no. 360742).

## DATA AVAILABILITY

The data and code for reproducing the results will be made openly available upon acceptance of the manuscript.

