## Supplementary material for "Can evolutionary potential help plants survive climate change? High variability in evolvability and G-matrix structure of *Hypericum* populations": Table S2; Table S3

| Gmatrix | Comparison type | Compared populations | cor (posterior medians) | cor (posterior means) | correlations of pair-wise samples | | | |
|---|---|---|---|---|---|---|---|---|
| | | | | | median _cor | mean_ cor | Lower CI | Upper CI |
| $\boldsymbol{G}_{F+V}$ | Interspecific | Hmac_Pooled-Hmon_Pooled | 0.371 | 0.274 | 0.16 | 0.178 | -0.184 | 0.641 |
| $\boldsymbol{G}_{F+V}$ | Intraspecific | Hmac_MHH5-Hmac_MH1 | 0.467 | 0.499 | 0.207 | 0.212 | -0.248 | 0.705 |
| $\boldsymbol{G}_{F+V}$ | Intraspecific | Hmac_MHH5-Hmac_Mei | 0.788 | 0.736 | 0.321 | 0.319 | -0.191 | 0.770 |
| $\boldsymbol{G}_{F+V}$ | Intraspecific | Hmac_MH1-Hmac_Mei | 0.154 | 0.227 | 0.064 | 0.062 | -0.269 | 0.391 |
| $\boldsymbol{G}_{ssF}$ | Interspecific | Hmac_Pooled-Hmon_Pooled | 0.542 | 0.825 | 0.146 | 0.142 | -0.750 | 0.930 |
| $\boldsymbol{G}_{ssF}$ | Intraspecific | Hmac_MHH5-Hmac_MH1 | 0.801 | 0.816 | 0.157 | 0.151 | -0.720 | 0.926 |
| $\boldsymbol{G}_{ssF}$ | Intraspecific | Hmac_MHH5-Hmac_Mei | 0.571 | 0.846 | 0.224 | 0.189 | -0.652 | 0.905 |
| $\boldsymbol{G}_{ssF}$ | Intraspecific | Hmac_MH1-Hmac_Mei | 0.925 | 0.974 | 0.344 | 0.291 | -0.567 | 0.938 |
| $\boldsymbol{G}_{ssV}$ | Interspecific | Hmac_Pooled-Hmon_Pooled | -0.407 | -0.402 | -0.221 | -0.162 | -0.746 | 0.735 |
| $\boldsymbol{G}_{ssV}$ | Intraspecific | Hmac_MHH5-Hmac_MH1 | 0.124 | 0.148 | 0.035 | 0.018 | -0.763 | 0.749 |
| $\boldsymbol{G}_{ssV}$ | Intraspecific | Hmac_MHH5-Hmac_Mei | 0.863 | 0.776 | 0.419 | 0.360 | -0.499 | 0.938 |
| $\boldsymbol{G}_{ssV}$ | Intraspecific | Hmac_MH1-Hmac_Mei | -0.168 | -0.154 | -0.131 | -0.122 | -0.635 | 0.442 |
| $\boldsymbol{G}_{V}$ | Interspecific | Hmac_Pooled-Hmon_Pooled | 0.516 | 0.484 | 0.338 | 0.294 | -0.446 | 0.866 |
| $\boldsymbol{G}_{V}$ | Intraspecific | Hmac_MHH5-Hmac_MH1 | -0.017 | -0.008 | -0.026 | -0.003 | -0.602 | 0.686 |
| $\boldsymbol{G}_{V}$ | Intraspecific | Hmac_MHH5-Hmac_Mei | 0.861 | 0.861 | 0.737 | 0.676 | 0.036 | 0.965 |
| $\boldsymbol{G}_{V}$ | Intraspecific | Hmac_MH1-Hmac_Mei | 0.350 | 0.332 | 0.314 | 0.290 | -0.298 | 0.807 |
| $\boldsymbol{G}_{V}$ | Intraspecific | Hmon_G-Hmon_Loh | 0.386 | 0.501 | 0.169 | 0.126 | -0.761 | 0.895 |
| $\boldsymbol{G}_{LH}$ | Interspecific | Hmac_Pooled-Hmon_Pooled | 0.555 | 0.458 | 0.299 | 0.268 | -0.351 | 0.776 |
| $\boldsymbol{G}_{LH}$ | Intraspecific | Hmac_MHH5-Hmac_MH1 | 0.791 | 0.791 | 0.556 | 0.492 | -0.205 | 0.928 |
| $\boldsymbol{G}_{LH}$ | Intraspecific | Hmac_MHH5-Hmac_Mei | 0.326 | 0.29 | 0.211 | 0.218 | -0.220 | 0.663 |
| $\boldsymbol{G}_{LH}$ | Intraspecific | Hmac_MH1-Hmac_Mei | 0.531 | 0.46 | 0.313 | 0.301 | -0.142 | 0.712 |

**Table S2.** Population pair-wise comparisons of G-matrices with random skewers. Two leftmost values give Pearson correlations of posterior distribution medians and means, and the four rightmost values give the median, mean and 95% confidence interval of Pearson correlations of posterior pair-wise samples.

| G-matrix | P-matrix | | | | | |
|---|---|---|---|---|---|---|
| $\boldsymbol{G}_{F+V}$ | Hmac | Hmon | Hmac_ MHH5 | Hmac_ MH1 | Hmac_ Mei | Hmon_ G |
| Hmac_Pooled | 0.773 | 0.058 | 0.748 | 0.672 | 0.447 | 0.058 |
| Hmac_MHH5 | 0.641 | 0.075 | 0.668 | 0.405 | 0.507 | 0.075 |
| Hmac_MH1 | 0.544 | -0.034 | 0.462 | 0.761 | 0.037 | -0.034 |
| Hmac_Mei | 0.670 | 0.207 | 0.568 | 0.263 | 0.831 | 0.207 |
| Hmon_G | 0.111 | 0.452 | -0.012 | 0.147 | 0.112 | 0.452 |

| $\boldsymbol{G}_{ssF}$ | Hmac | Hmon | Hmac_ MHH5 | Hmac_ MH1 | Hmac_ Mei | Hmon_ G |
|---|---|---|---|---|---|---|
| Hmac_Pooled | 0.770 | 0.883 | 0.859 | 0.900 | 0.433 | 0.883 |
| Hmac_MHH5 | 0.175 | 0.043 | 0.275 | 0.181 | 0.058 | 0.043 |
| Hmac_MH1 | 0.713 | 0.481 | 0.756 | 0.648 | 0.615 | 0.481 |
| Hmac_Mei | 0.879 | 0.766 | 0.911 | 0.875 | 0.702 | 0.766 |
| Hmon_G | -0.014 | 0.463 | 0.138 | 0.288 | -0.430 | 0.463 |

| $\boldsymbol{G}_{ssV}$ | Hmac | Hmon | Hmac_ MHH5 | Hmac_ MH1 | Hmac_ Mei | Hmon_ G |
|---|---|---|---|---|---|---|
| Hmac_Pooled | 0.999 | -0.254 | 0.922 | 0.810 | 0.454 | -0.254 |
| Hmac_MHH5 | 0.849 | -0.102 | 0.824 | 0.374 | 0.794 | -0.102 |
| Hmac_MH1 | 0.611 | -0.314 | 0.474 | 0.903 | -0.215 | -0.314 |
| Hmac_Mei | 0.584 | 0.137 | 0.458 | 0.072 | 0.939 | 0.137 |
| Hmon_G | -0.421 | 0.971 | -0.565 | -0.278 | -0.082 | 0.971 |

| $\boldsymbol{G}_{V}$ | Hmac | Hmon | Hmac_ MHH5 | Hmac_ MH1 | Hmac_ Mei | Hmon_ G | Hmon_ Loh |
|---|---|---|---|---|---|---|---|
| Hmac_Pooled | 0.924 | 0.407 | 0.731 | 0.659 | 0.746 | 0.378 | 0.411 |
| Hmon_Pooled | 0.418 | 0.984 | 0.237 | 0.033 | 0.648 | 0.935 | 0.986 |
| Hmac_MHH5 | 0.601 | 0.880 | 0.359 | 0.123 | 0.849 | 0.818 | 0.889 |
| Hmac_MH1 | 0.713 | -0.219 | 0.725 | 0.886 | 0.114 | -0.203 | -0.222 |
| Hmac_Mei | 0.760 | 0.574 | 0.511 | 0.301 | 0.899 | 0.554 | 0.572 |
| Hmon_G | 0.481 | 0.467 | 0.558 | 0.566 | 0.060 | 0.518 | 0.440 |
| Hmon_Loh | 0.132 | 0.958 | -0.074 | -0.193 | 0.496 | 0.988 | 0.930 |

| $\boldsymbol{G}_{LH}$ | Hmac | Hmon | Hmac_ MHH5 | Hmac_ MH1 | Hmac_ Mei | Hmon_ G |
|---|---|---|---|---|---|---|
| Hmac_Pooled | 0.922 | 0.452 | 0.826 | 0.954 | 0.503 | 0.452 |
| Hmac_MHH5 | 0.960 | 0.460 | 0.964 | 0.909 | 0.236 | 0.460 |
| Hmac_MH1 | 0.884 | 0.525 | 0.767 | 0.963 | 0.436 | 0.525 |
| Hmac_Mei | 0.437 | 0.110 | 0.271 | 0.449 | 0.985 | 0.110 |
| Hmon_G | 0.685 | 0.952 | 0.681 | 0.680 | 0.081 | 0.952 |

Table S3. Pairwise comparisons (Pearson correlations) of G-matrices and the respective P- matrices through 1000 simulated values of evolvability along random selection gradients given the derived G- and P-matrices of studied *H. maculatum* and *H. montanum* populations. The two leftmost columns show comparisons of G with species-specific P-matrices (mean P over populations), while the remaining columns show comparisons with population-specific P-matrices. Shaded cells show correlations of the respective population- and species-specific G- and P-matrices.
